# Complete sequencing of a cynomolgus macaque major histocompatibility complex haplotype

**DOI:** 10.1101/2022.10.24.513544

**Authors:** Julie A. Karl, Trent M. Prall, Hailey E. Bussan, Joshua M. Varghese, Aparna Pal, Roger W. Wiseman, David H. O’Connor

**Author notes:** These authors contributed equally to this work (Julie A. Karl, Trent M. Prall). Corresponding author: David H. O’Connor.

## Abstract

Macaques provide the most widely used nonhuman primate models for studying immunology and pathogenesis of human diseases. While the macaque major histocompatibility complex (MHC) region shares most features with the human leukocyte antigen (HLA) region, macaques have an expanded repertoire of MHC class I genes. Although a chimera of two rhesus macaque MHC haplotypes was first published in 2004, the structural diversity of MHC genomic organization in macaques remains poorly understood due to a lack of adequate genomic reference sequences. We used ultra-long Oxford Nanopore and high-accuracy PacBio HiFi sequences to fully assemble the ∼5.2 Mb M3 haplotype of an MHC-homozygous, Mauritian-origin cynomolgus macaque (*Macaca fascicularis*). The MHC homozygosity allowed us to assemble a single MHC haplotype unambiguously and avoid chimeric assemblies that hampered previous efforts to characterize this exceptionally complex genomic region in macaques. The high quality of this new assembly is exemplified by the identification of an extended cluster of six *Mafa-AG* genes that contains a recent duplication with a remarkably similar ∼48.5 kb block of sequence. The MHC class II region of this M3 haplotype is similar to the previously sequenced rhesus macaque haplotype and HLA class II haplotypes. The MHC class I region, in contrast, contains 13 *MHC-B* genes, four *MHC-A* genes, and three *MHC-E* genes (versus 19 *MHC-B*, two *MHC-A*, and one *MHC-E* in the previously sequenced haplotype). These results provide an unambiguously assembled single contiguous cynomolgus macaque MHC haplotype with fully curated gene annotations that will inform infectious disease and transplantation research.

## Introduction

Rhesus (*Macaca mulatta*) and cynomolgus (*Macaca fascicularis*) macaques are the most widely-used nonhuman primate models for infectious disease and transplantation research (Anderson and Kirk 2013; Burwitz et al. 2017; Knechtle et al. 2019; Shiina and Blancher 2019; D’Souza et al. 2022). Proper utilization of a model organism of human disease requires a deep understanding of the immune system and the genetic determinants that govern its responses. The major histocompatibility complex (MHC) encodes an array of gene products that are critical for host immune responses against infection as well as transplanted cells and tissues (Horton et al. 2004; Trowsdale and Knight 2013; Shiina et al. 2017). The most widely studied of these are the MHC class I (MHC-I) proteins that contain highly polymorphic peptide binding domains that play key roles in adaptive immunity by displaying endogenous antigens on the cell surface for surveillance by CD8+ cytotoxic T cells. Many MHC-I proteins also play an important role in innate immunity by serving as ligands for killer-cell immunoglobulin-like receptors (KIR) on natural killer cells (NK) (Trowsdale and Knight 2013). In contrast, MHC class II (MHC-II) proteins form heterodimers to present exogenous peptides to CD4+ helper T cells. MHC-II expression is largely confined to a subset of lymphoid cell types while MHC-I proteins are expressed by most tissues. Several additional genes in the MHC-II region encode proteins involved in proteasome processing (*PSMB8, PSMB9*) and peptide antigen transport (*TAP1, TAP2, TAPBP*) (Horton et al. 2004). The MHC class III genomic region, which lies between MHC-I and MHC-II regions, also encodes several classes of immune proteins including cytokines (*TNF, LTA, LTB*) and the complement cascade (*C2, C4A, C4B, CFB*) (Horton et al. 2004).

Genome-wide association studies have implicated more associations between the MHC and human autoimmune, inflammatory and infectious diseases than any other region of the genome (Horton et al. 2004; Trowsdale and Knight 2013). Although many of these human disease associations have been attributed to specific MHC-I and MHC-II alleles, linkage disequilibrium in this gene-dense region makes identification of specific causative variants for disease phenotypes extremely challenging. Non-immune system loci that make up the majority of genes in the MHC region are also likely to be responsible for a number of disease associations, so characterizing allelic variation in these genes is also essential for establishing genotype/phenotype correlations.

Unfortunately, the reference genome sequences for the MHC of macaques remain inaccurate, especially in the MHC class I region. Unlike human leukocyte antigen (HLA) haplotypes which contain three highly polymorphic class I genes (*HLA-A, HLA-B*, and *HLA-C*), the macaque MHC genomic region has undergone complex duplication, deletion, and rearrangement events such that different haplotypes contain a variable number of class I loci (Daza-Vamenta et al. 2004; Fukami-Kobayashi et al. 2005; Shiina et al. 2006; Watanabe et al. 2007). Sequencing of an MHC class I B haplotype from a rhesus macaque bacterial artificial chromosome (BAC) library (CHORI-250) identified 19 distinct *MHC-B* genes (Daza-Vamenta et al. 2004; Shiina et al. 2006). Extensive sequencing of MHC class I transcript and genomic DNA PCR amplicons suggests that this dramatic expansion of *MHC-B* genes is common in both rhesus and cynomolgus macaques (Otting et al. 2005; Doxiadis et al. 2013; Wiseman et al. 2013; Karl et al. 2017; Shortreed et al. 2020). These *MHC-B* duplication events appear to have arisen after the divergence of macaques and humans from their most recent common ancestor approximately 27 million years (Myr) ago (Fukami-Kobayashi et al. 2005). Macaques lack a clear orthologue of *HLA-C* that is estimated to have been duplicated from *HLA-B* approximately 22 Myr ago (Fukami-Kobayashi et al. 2005). Furthermore, transcript sequencing has provided evidence for the existence of multiple *MHC-A* and *MHC-E* genes per individual macaque haplotype (Wu et al. 2018; de Groot et al. 2020). In contrast to humans, macaques have lost a functional nonclassical *MHC-G* gene. The *HLA-G* locus has been replaced by a fusion of *MHC-A* and *MHC-G* genes, referred to as *MHC-AG* that is expressed in placenta and contributes to maternal tolerance of fetuses (Boyson et al. 1999). The *MHC-AG* gene regions of both rhesus and cynomolgus macaques have undergone a similar duplication as their classical MHC class I counterparts *MHC-A* and *MHC-B* (Daza-Vamenta et al. 2004; Shiina et al. 2006, 2017).

The implications for macaque MHC class I gene duplication are unclear. It has been established that various *MHC-A* and *MHC-B* genes exhibit differential levels of steady state RNA that we distinguish with ‘major’ and ‘minor’ allele designations (Otting et al. 2005; Budde et al. 2010; Karl et al. 2013).

Relative transcriptional levels of MHC class I genes result in differential cell surface expression (Rosner et al. 2010). Several rhesus macaque MHC class I molecules have been shown to mainly exist in subcellular compartments suggesting functional diversification of duplicated genes (Rosner et al. 2010). In contrast to humans, particular macaque MHC class I genes are expressed differentially in distinct leukocyte subsets suggesting the existence of differential function amongst duplicated genes (Greene et al. 2011). Furthermore, MHC expression is known to vary wildly between tissue subsets Boegel et al. 2018). Genomic assemblies of multiple MHC region configurations are a prerequisite to understanding the differential functions of these duplicated genes and how macaques manage to balance an expanded repertoire of MHC class I genes without causing autoimmunity.

Due to the importance of macaques in immunological research, thousands of classical and nonclassical class I and class II macaque MHC cDNA transcript sequences have been cataloged (Shiina and Blancher 2019). However, the genomic context of these transcript sequences is largely unexplored. A single Indian rhesus macaque MHC haplotype was sequenced by Daza-Vamenta *et al*. in 2004 and resequenced by Shiina *et al*. in 2006 (Daza-Vamenta et al. 2004; Shiina et al. 2006). In both cases, the overlapping BACs were derived from an MHC-heterozygous macaque, leading to chimeric sequences that do not reflect a single, intact MHC haplotype. Obtaining additional MHC haplotype sequences has been complicated by the high degree of conservation among the closely related, duplicated genes. This difficulty is demonstrated by gaps and artificial sequence mergers in the MHC class I regions of publicly available rhesus (Mmul_10) and cynomolgus (MFA1912RKSv2) macaque genome reference builds (Warren et al. 2020; Jayakumar et al. 2021).

Recently, Nurk and colleagues reported the first *de novo* sequencing of a complete human genome using ultra-long Oxford Nanopore Technologies (ONT) sequencing reads (Nurk et al. 2022). This human genome build (T2T-CHM13) includes five new chromosome arms representing an additional 8% of the genome that had not been previously sequenced due to technical limitations. Nanopore sequencing enables intact DNA fragments as long as 4 Megabase pairs (Mb) to be sequenced in their entirety (Genome assembly 2022). This allows repetitive genomic blocks to be bridged by unambiguously assembling ultra-long DNA reads into chromosome-level scaffolds (Jain et al. 2018; Nurk et al. 2022). Similar repetitive sequence motifs such as transposable elements, short tandem repeats, and homopolymer stretches are abundant within the macaque MHC. Therefore, applying state-of-the-art sequencing techniques to this region will result in an unprecedented resolution of the genomic architecture. While individual ONT sequencing reads have a higher per-base error rate than other next-generation sequencing platforms, the ultra-long scaffolds can be combined with higher-accuracy sequencing for error correction. Traditionally researchers used Illumina short-read sequencing for this purpose, but recent improvements in Pacific Biosciences (PacBio) methods for HiFi circular consensus sequencing routinely produce high-accuracy reads of >10 kilobase pairs (kb) (Wenger et al. 2019; Nurk et al. 2020). This provides improved error correction versus short reads through increased confidence in unique mapping across the scaffold assembly with an ability to distinguish between even repetitive sequence elements. Researchers are beginning to use PacBio HiFi sequencing for gap filling and sequence polishing of ONT scaffolds to create unprecedentedly complete high-quality assemblies (Kakuk et al. 2021; Nurk et al. 2022; Wang et al. 2022).

The genetic diversity of Mauritian-origin cynomolgus macaques (MCM) is far less than that of other species of macaques with research significance due to geographic isolation and a founder effect (Sussman and Tattersall 1986). The entire MHC diversity within the geographic population can be characterized by seven ancestral haplotypes (Wiseman et al. 2013). This unique genetic homogeneity allows for the selection of MHC-homozygous individuals with relative ease. For the current study we sequenced an animal that is homozygous for the M3 MHC haplotype. This M3 haplotype is the second most abundant ancestral haplotype in the MCM population, accounting for 14% of the MHC haplotypes overall (Wiseman et al. 2013). Furthermore, cDNA allelic transcripts are fully characterized for most of the variable MHC classical class I and class II genes, allowing us to estimate the error rate of genomic assemblies accurately. Here we take advantage of the strengths of ONT and PacBio HiFi sequencing to generate a ∼60x ONT and ∼30x PacBio HiFi coverage genome of an MHC-homozygous MCM. Following assembly and error correction of the full genomic MHC region, we obtained a ∼5.2 Mb contig spanning the MHC class I, class II, and class III regions, creating the most complete macaque MHC sequence to date. We found structural rearrangements in the *MHC-B* region relative to the only previously sequenced macaque *MHC-B* haplotype, resulting in 13 *MHC-B* genes. We also found an expansion of *MHC-A, MHC-AG*, and *MHC-E* genes on this haplotype, underscoring that *MHC-B* is not the only locus in the MHC exhibiting copy number variation in macaques. This full genomic MHC region assembly provides an important new comparator for understanding differences between macaque and human MHC and, in turn, cellular immunity. It establishes a new paradigm for sequencing macaque MHC haplotypes that will likely lead to the solution of many similar haplotypes in the future.

## Results

### Mauritian-origin cynomolgus macaque M3 haplotype gene content

The complete gene content of the ∼5.2 Mb cy0333 genomic region MHC M3 haplotype of MCM is displayed in **Fig. 1**. This represents the first detailed assembly and annotation focused solely on the MHC genomic region of a macaque in almost two decades since both Daza-Vamenta *et al*. and Shiina *et al*. characterized the MHC region of a rhesus macaque BAC library (CHORI-250) (Daza-Vamenta et al. 2004; Shiina et al. 2006). The cost and labor required to generate and sequence BAC libraries with older sequencing technologies made such studies into this complex and immune-important region prohibitively expensive. The unique limited diversity of the MCM population also enabled the selection of an MHC-homozygous sample, ensuring that the resulting assembly would unambiguously represent a single contiguous macaque MHC haplotype. A total of 383 putative genes orthologous to either human or rhesus macaque MHC genes were identified on the cy0333 M3 haplotype (**Supplemental Table S1**). Of these, 152 contain open reading frames indicative of potential protein expression. This is roughly equivalent to the 305 genes (145 protein-coding) on the human reference genome GRCh38. While a pair of annotated cynomolgus macaque reference genomes are also currently available (Macaca_fascicularis_5.0; GCF_000364345 and MFA1912RKSv2; GCF_012559485), they were not used to determine gene content of the cy0333 MHC M3 haplotype. The current annotation builds for cynomolgus macaques are not as thorough as human and rhesus macaque MHC region annotations since there are many uncharacterized loci which are difficult to compare to the carefully curated human gene annotations.

**Figure 1.**
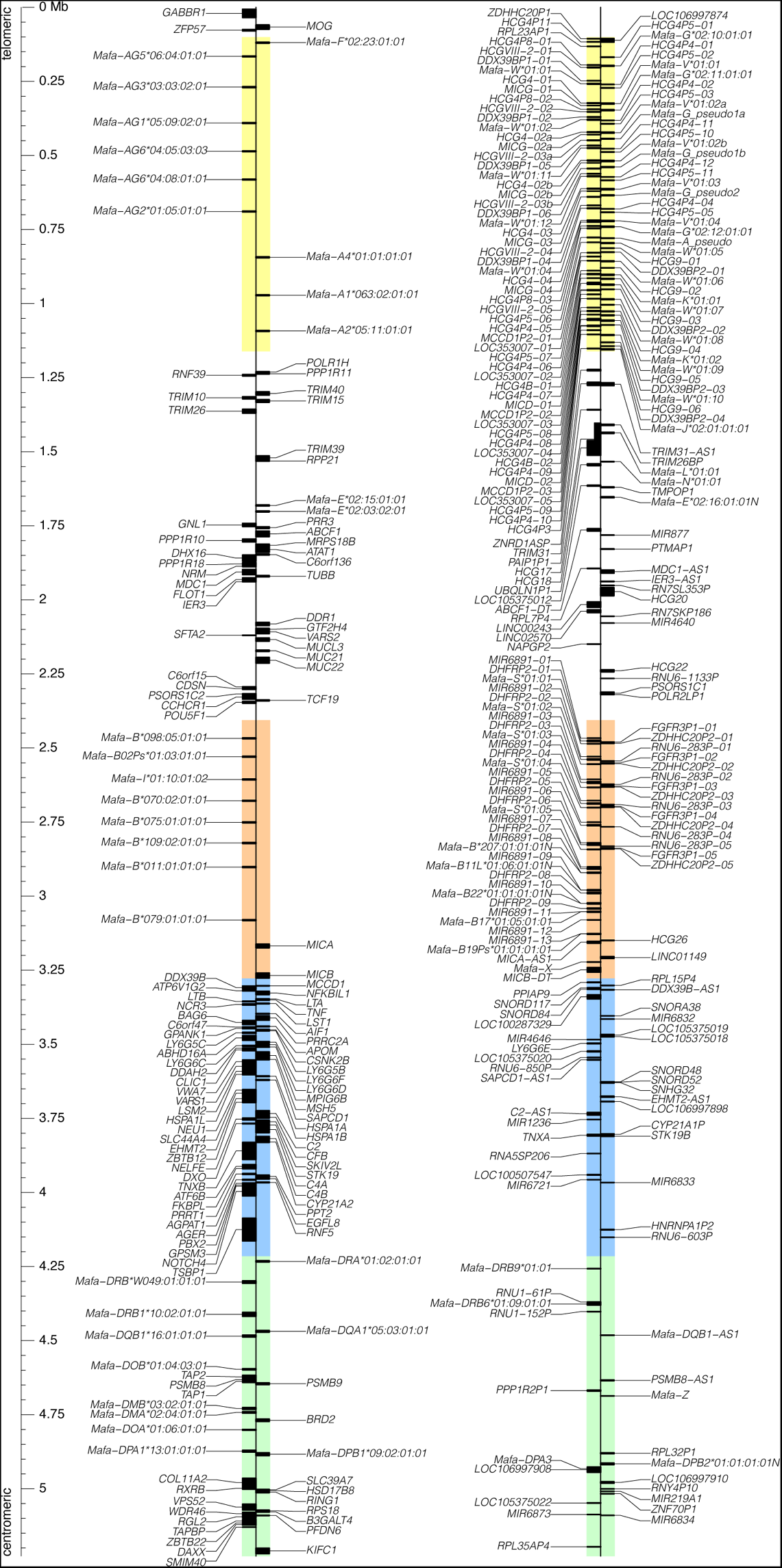
Gene content of the cy0333 MHC M3 haplotype of MCM full genomic region. The MHC full genomic region is defined as everything from the telomeric gene *GABBR1* to the centromeric gene *KIFC1*, as per Shiina *et al*. (Shiina et al. 2017). Gene content is separated into putatively expressed genes on the left line and non-coding RNAs and pseudogenes on the right line. Gene sequences associated with the telomeric to centromeric sense strand are displayed to the right of each line, and those associated with the anti-sense strand are displayed to the left. The class I A region is highlighted in yellow, the class I B region is highlighted in orange, the class III region is highlighted in blue, and the class II region is highlighted in green. The position in Mb is shown on the left.

Segmental duplications have significantly increased the gene content of the cy0333 M3 haplotype of MCM within the MHC class I A and class I B regions compared to the human reference genome (**Fig. 2**). The cy0333 haplotype contains four total *Mafa-A* genes, three of which have canonical classical MHC class I gene open reading frames, and one pseudogene in which the first ∼619 nucleotides of the expected open reading frame (all of exons 1-3) are missing due to a large deletion event. The *Mafa-B* region experienced even more duplication events than the *Mafa-A* region, with a total of 13 *Mafa-B* genes. Eight of these contain canonical classical MHC class I B gene open reading frames, but the other five cannot be translated into traditional MHC class I B proteins due to the presence of premature stop codons or frame shifts from the canonical class I B open reading frame.

**Figure 2.**
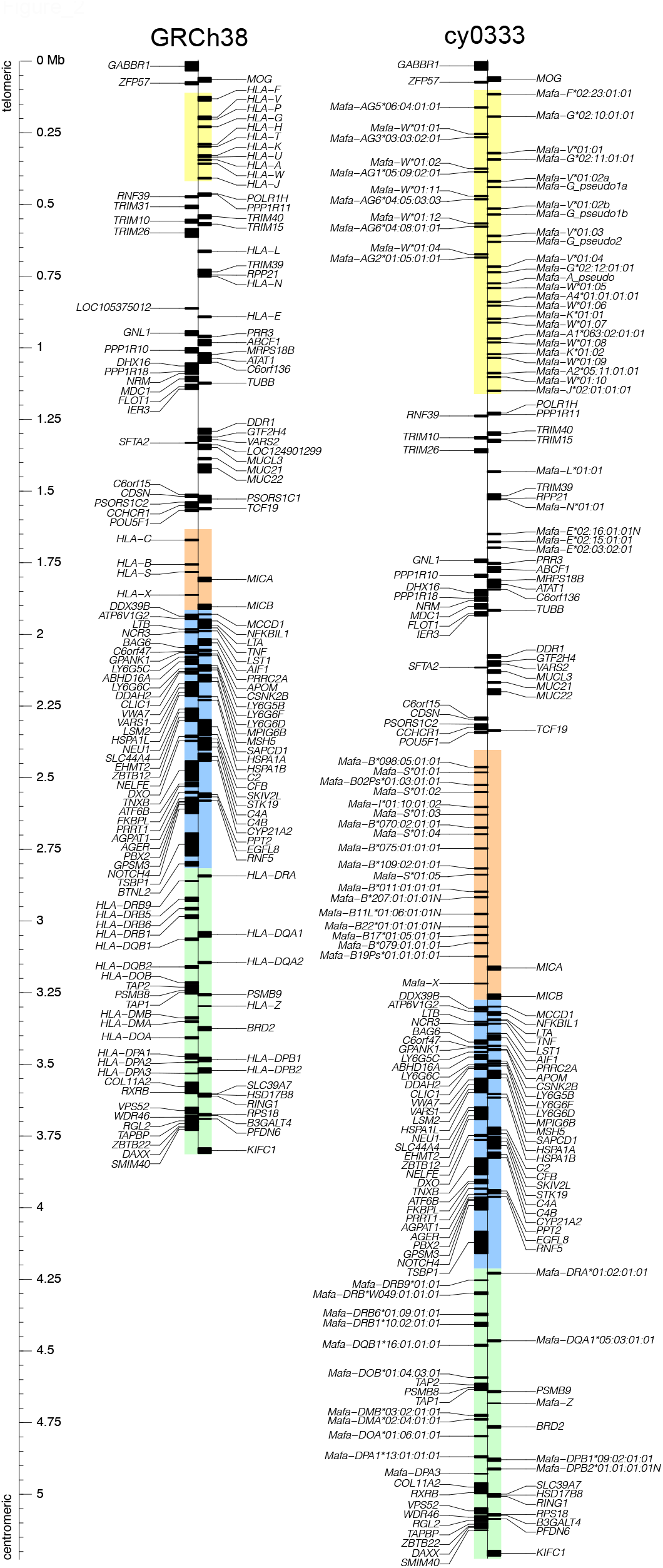
Comparison of the representative gene content of the human and the cy0333 MCM full genomic MHC regions. Protein coding genes and select nonclassical MHC class I and class II pseudogenes are shown for the human genome reference GRCh38 (left) and cy0333 (right). The class I A region is highlighted in yellow, the class I B region is highlighted in orange, the class III region is highlighted in blue, and the class II region is highlighted in green. Both sequences are equivalently scaled and presented from telomeric genes at the top to centromeric genes at the bottom. Gene sequences associated with the telomeric to centromeric sense strand are displayed to the right of each line, and those associated with the anti-sense strand are displayed to the left. The relative position in Mb for both sequences is shown on the left. Significant duplication of genes, particularly in the class I A and class I B regions, contribute to the extra ∼1.4 Mb of sequence in cy0333.

In addition to differences in classical class I gene content between humans and macaques, there are differences in nonclassical class I genes, namely the *MHC-G* and *MHC-AG* genes. In humans, the *HLA-G* gene is functional; in macaques, all copies of the *MHC-G* gene are pseudogenes with early stop codons introduced in exon 3 (some *MHC-G* genes per macaque haplotype may also lack traditional start sequences – the cy0333 MHC M3 haplotype of MCM contains three such *Mafa-G* genes). Macaques have instead gained putatively expressed *MHC-AG* genes, which appear to be a combination of the 5’ end of an *MHC-A* gene (which recovers the stop codon in exon 3) plus the 3’ end of an *MHC-G* gene (which maintains the relatively early stop codon in exon 6 that is also observed in the functional *HLA-G* gene). The MHC M3 haplotype of MCM contains six distinct *Mafa-AG* genes and six duplications of the *Mamu-G* gene. The MCM M3 haplotype also has three total nonclassical class I *Mafa-E* sequences (two expressed *Mafa-E* genes plus one non-expressed pseudogene *Mafa-E*02:16:01:01N* lacking a traditional start codon and containing an early stop codon in exon 3). This differing number of segmental duplications is typical for macaques – different haplotypes have undergone different numbers of segmental duplications of classical and nonclassical genes in the class I regions (Daza-Vamenta et al. 2004; Fukami-Kobayashi et al. 2005; Shiina et al. 2006; Otting et al. 2007; Watanabe et al. 2007; Karl et al. 2013; Wiseman et al. 2013; Shortreed et al. 2020).

These segmental duplication events account for most of the ∼1.4 Mb length difference between human reference GRCh38 and MCM cy0333; the remaining MHC class III and class II regions are relatively well-conserved (**Fig. 2**). The most significant differences in these regions are in the *MHC-DRB* genes, where not even all human haplotypes contain the same numbers of expressed and non-expressed *HLA-DRB* genes.

All but five of the putatively expressed MHC genes in the human reference genome (*LOC124901299, HLA-C, BTNL2, HLA-DQA2*, and *HLA-DQB2*) appear to have orthologs on the MHC M3 haplotype of MCM. There are more differences in the content of non-coding RNAs and pseudogenes between the two species, with 56 human non-coding genes appearing to lack a cynomolgus macaque homolog. These pseudogenes may have been lost or gained after humans and macaques diverged.

### Comparison to the MHC region of other macaque reference genomes

We believe that our genomic region assembly of the MHC M3 haplotype of MCM is the most contiguous macaque MHC region assembly generated to date – there are no gaps or unknown N sequence fillers across the entire 5,227,476 base pairs (bp). With the advent of next-generation sequencing more than a decade ago, it became faster and less expensive to sequence whole genomes. This led to many new whole genome assemblies from Illumina short-read sequence data across multiple model organisms, like one of the representative cynomolgus macaque genomes Macaca_fascicularis_5.0 (GCF_000364345). This genome was assembled from Illumina HiSeq shotgun sequencing reads from a female Indonesian-origin cynomolgus macaque at ∼68x genome coverage in 2013 (https://www.ncbi.nlm.nih.gov/assembly/GCF_000364345.1/). While overall advantageous due to its relatively low cost and reasonably accurate assembly in non-repetitive regions, these short reads are insufficient to assemble regions with segmental duplications like the MHC accurately. This is quite apparent when comparing the alignment of our cy0333 MHC M3 haplotype of MCM to the Macaca_ fascicularis_5.0 assembly for those regions (**Fig. 3**). The white bars within the Macaca_fascicularis_5.0 reference correspond to gaps of unknown sequence indicated with Ns in the reference sequence. There are 230 total gaps covering at least 373,286 total bp out of the 3.9 Mb shown for Macaca_ fascicularis_5.0, so over 9.5% of the reported MHC sequence for this reference genome is unknown.

**Figure 3.**
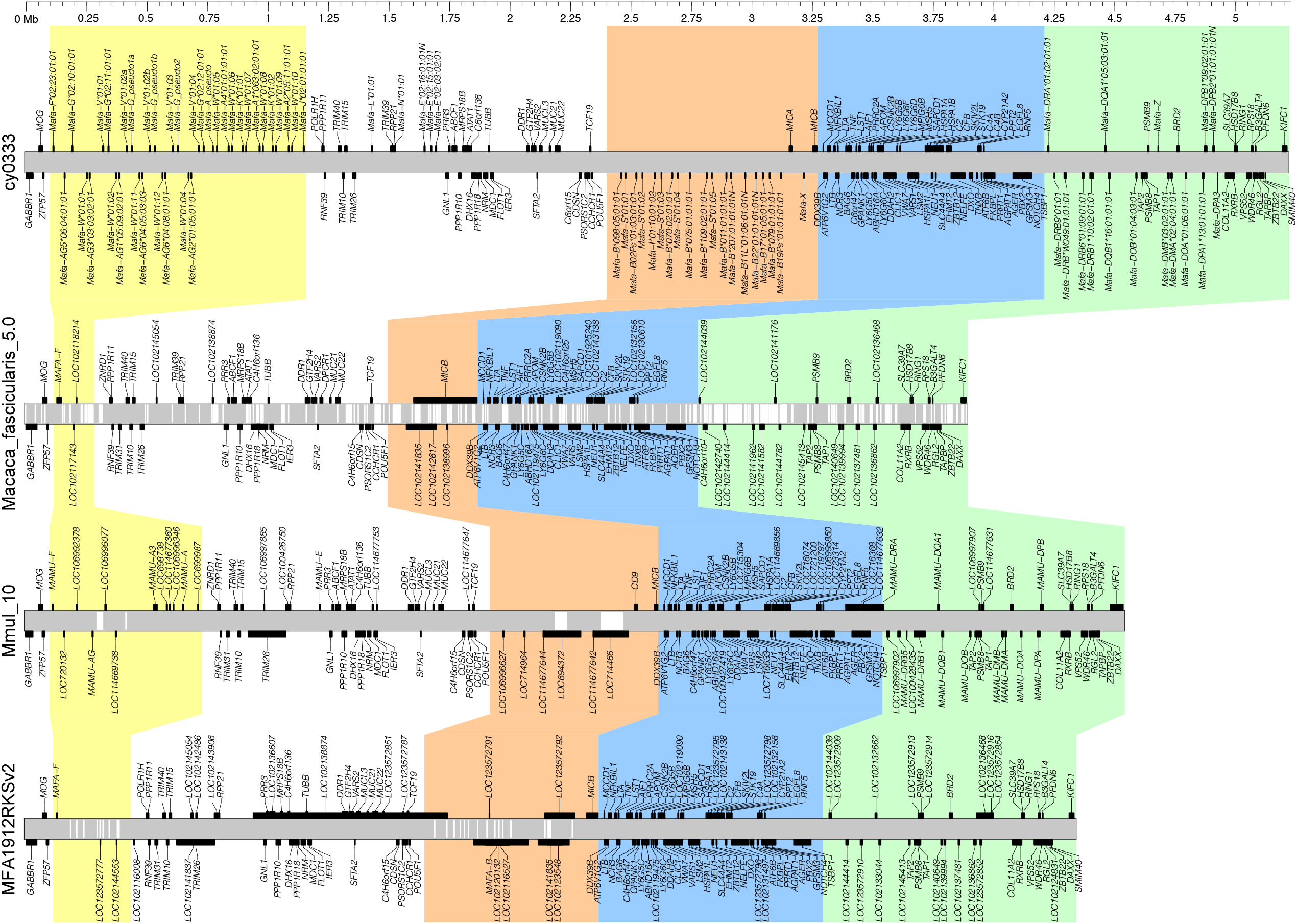
Continuity of the cy0333 MCM full genomic MHC region versus three publicly available macaque reference genomes. The cy0333 MHC region (OP204634) is shown in the top panel, the NCBI cynomolgus macaque reference genome Macaca_fascicularis_5.0 (GCF_000364345) assembled from Illumina short-read data is shown in the second panel down, the rhesus macaque reference genome Mmul_10 (GCF_003339765) assembled from ∼13.5 kb PacBio RSII data is shown in the third panel down, and the NCBI cynomolgus macaque reference genome MFA1912RKSv2 (GCF_012559485) assembled from ∼12 kb PacBio Sequel II data is shown in the bottom panel. All sequences are uniformly scaled, and the position in Mb for all sequences is shown across the top. Gray tracks represent the sequence for each reference, and white gaps within the gray tracks represent gaps in the assembly. The class I A region is highlighted in yellow, the class I B region is highlighted in orange, the class III region is highlighted in blue, and the class II region is highlighted in green.

The full genomic MHC region of Macaca_fascicularis_5.0 is also notably over 1.3 Mb shorter than cy0333 and, in particular, lacks gene density within the MHC class I A and class I B regions. There is no reason to believe that the Indonesian cynomolgus macaque sequenced for the Macaca_ fascicularis_5.0 assembly would not have a similar MHC class I gene density as the cy0333 M3 haplotype of MCM. Mauritian cynomolgus macaques are most likely derived from a small population of Indonesian cynomolgus macaques, and previous MHC haplotyping studies of cynomolgus macaques from Indonesia as well as other geographic origins have shown expected levels of segmental duplications for these regions (Shortreed et al. 2020; de Groot et al. 2022). However, the MHC class I A and class I B regions for Macaca_fascicularis_5.0 are both significantly shorter than their cy0333 counterparts.

With the difficulties inherent to the accurate assembly of highly repetitive regions, efforts are underway to improve reference genome assemblies with alternative next-generation sequencing technologies like PacBio. These technologies are currently more expensive per base pair than Illumina sequencing but can produce longer reads to assist in the assembly of repetitive regions. The rhesus macaque reference genome released in 2019, Mmul_10 (GCF_003339765), was assembled from PacBio RSII sequencing reads from a female Indian rhesus macaque fibroblast cell line at ∼66x genome coverage (https://www.ncbi.nlm.nih.gov/assembly/GCF_003339765.1/) (Warren et al. 2020). With average sequence lengths much longer than Illumina at ∼13.5 kb, this technology is superior though still not perfect for the MHC region. Alignment of the MHC region from the Mmul_10 reference genome to the MCM M3 haplotype shows that it is only ∼677 kb shorter than its cy0333 counterpart (**Fig. 3**). Mmul_10 contains expanded MHC class I A and class I B regions compared to Macaca_ fascicularis_5.0 though each region is still shorter than the equivalent region in cy0333. Mmul_10 also notably only has four gaps of Ns – two gaps within the class I A region of 22,370 bp and 5 bp as well as two gaps of 45,930 bp and 86,789 bp within the class I B region (∼3.4% of the reported 4.5 Mb of the Mmul_10 reference MHC full genomic region).

A quality assessment of the Mmul_10 MHC region showed that the AG07107 fibroblast line used for this assembly was homozygous for the Mamu-A004 haplotype (the most common MHC class I A haplotype in rhesus macaques) (Warren et al. 2020). The lack of heterozygosity for the genomic region encoding the two functional *Mamu-A* genes of this haplotype made it possible to assemble accurately, harnessing the same advantage we had in assembling the fully homozygous M3 MHC genomic region of cy0333. This localized area of homozygosity resulted in an area of the MHC class I A region that contained no unknown sequence gaps and a relatively clean annotation despite the segmentally duplicated *Mamu-A* genes. The ∼22 kb of gaps within the MHC class I A region of Mmul_10 are near the *Mamu-AG* genes, indicating some uncertainty about the accuracy of the assembly in that portion of the class I A region. The MHC class I B region of Mmul_10 also includes two relatively large gaps. As presented in the Mmul_10 primary chromosome 4 assembly, this region is a fusion of the two MHC class I B haplotypes (Mamu-B048 and Mamu-B055) from the heterozygous AG07107 fibroblasts (Warren et al. 2020). It contains four of the genes expected for the Mamu-B048 haplotype and two of the genes expected on the Mamu-B055 haplotype. Most of the remaining *Mamu-B* genes expected for the Mamu-B048 and Mamu-B055 haplotypes map to several unlocalized genomic scaffolds associated with the Mmul_10 whole genome assembly. While the MHC class II region is appropriately sized, the gene content again represents a mixture of the two MHC class II haplotypes in AG07107 fibroblast line, with several of the remaining genes from the alternate haplotype located on additional unlocalized genomic scaffolds.

Another cynomolgus macaque reference genome (MFA1912RKSv2; GCF_012559485) was assembled from PacBio Sequel II data recently. This assembly was generated from a Cambodian-origin cynomolgus macaque with ∼81x genome coverage and sequence read lengths averaging ∼12 kb (https://www.ncbi.nlm.nih.gov/data-hub/genome/GCF_012559485.2/) (Jayakumar et al. 2021). Comparison to our cy0333 MHC M3 haplotype again highlights a reasonable assembly across areas of less extreme segmental duplications like the MHC class III and class II regions as well as the MHC class I interval between the class I A and class I B regions (**Fig. 3**). However, there are once again gaps of Ns within the class I A (nine total gaps) and the class I B (16 total gaps) regions. These gaps are all shown as ∼500 bp; lack of variability in gap lengths indicates an extra degree of uncertainty to those gap regions – not only are the precise nucleotides unknown but so are the actual expected gap sizes. Overall, MFA1912RKSv2 contains 12,496 Ns across 4.35 Mb of the MHC full genomic region (∼0.3%), but this is likely an underestimate. There are also notable issues with the current annotations for MFA1912RKSv2, with no annotated *Mafa-A* genes, only a few *Mafa-B* genes where the predicted translations extend across several of the duplicated *Mafa-B* genes, and an ∼800 kb predicted translation for the *Mafa-E* gene *LOC102138874* (along with more appropriately-sized predicted translations for that same gene annotation).

Overall, longer reads are easier to assemble accurately, but heterozygosity still complicates *de novo* assemblies, especially for regions of highly duplicated genes like the MHC class I A and class I B regions. Our hybrid ONT and PacBio HiFi assembly for cy0333 (OP204634) provides a complete and fully annotated alternative genomic reference for the MHC region.

### Classical and nonclassical MHC class I and class II gene content

Previous cDNA studies of MCMs provided a roadmap for expected MHC classical class I and class II gene content for the M3 haplotype (Budde et al. 2010; O’Connor et al. 2010; Wiseman et al. 2013). In those studies, ten classical class I genes (three *Mafa-A* genes and seven *Mafa-B/-I* genes) and seven classical class II genes (two *Mafa-DRB* genes and one gene per locus for *Mafa-DRA, Mafa-DQA1, Mafa-DQB1, Mafa-DPA1*, and *Mafa-DPB1*) were identified. In this study, we detected those 17 expected M3 genes plus seven additional classical class I genes (one *Mafa-A* gene and six *Mafa-B* genes) and five more classical class II genes (two *Mafa-DRB* genes and one gene per locus for *Mafa-DQB1-AS1, Mafa-DPA3*, and *Mafa-DPB2*) (**Supplemental Table S2**). All but two of these additional 12 classical class I and class II genes lack canonical open reading frames and cannot be translated into traditional class I and class II proteins; one of the 12 is a non-protein coding anti-sense RNA of the traditional *Mafa-DQB1* locus (*Mafa-DQB1-AS1*). The remaining classical class I gene not previously identified on the MCM M3 haplotype, *Mafa-B02Ps*01:03:01:01*, does have a canonical seven-exon *Mafa-B* gene open reading frame but appears to be expressed poorly, if at all, in whole blood.

Beyond the classical class I and class II genes, this study also detected the full complement of 19 nonclassical class I and class II genes of the MHC M3 haplotype of MCM – one *Mafa-F* gene, six *Mafa-AG* genes, five distinct *Mafa-G* pseudogenes, three *Mafa-E* genes (one with a premature stop codon), and one gene each for the *Mafa-DMA, Mafa-DMB, Mafa-DOA*, and *Mafa-DOB* loci as well as 27 total distinct MHC region class I pseudogenes (one *Mafa-J* pseudogene, two *Mafa-K* pseudogenes, one *Mafa-L* pseudogene, one *Mafa-N* pseudogene, five *Mafa-S* pseudogenes, four *Mafa-V* pseudogenes, 11 *Mafa-W* pseudogenes, one *Mafa-X* pseudogene, and one *Mafa-Z* pseudogene) (**Supplemental Table S2**).

Three of the class I genes identified on the cy0333 MHC M3 haplotype of MCM, labeled *Mafa-A_ pseudo, Mafa-G_pseudo1*, and *Mafa-G_pseudo2*, lack official ImmunoPolymorphism Non-Human Primates MHC Database (IPD-MHC NHP) nomenclature because they are peculiar cases. These three pseudogenes are all missing the traditional start codon site due to large deletions – *Mafa-A_pseudo* is missing essentially all of exon 1 through intron 3, and both of the *Mafa-G* pseudogenes are missing exon 1 through most of exon 2 (only retaining ∼37 bp of exon 2) – indicating either incomplete duplication of those sequence blocks or large deletion events post-duplication. This is in contrast to the other duplicated class I genes lacking traditional open reading frames due to premature stop codons from internal single nucleotide polymorphisms or frameshifts within genes with typical start codon patterns (the other three *Mafa-G* pseudogenes, *Mafa-E*02:16:01:01N*, and the five *Mafa-B* pseudogenes).

We assessed the nucleotide accuracy of the cy0333 hybrid ONT and PacBio MHC M3 haplotype assembly with independent PacBio Sequel II HiFi sequencing of ∼2.2-5.0 kb gDNA amplicons spanning the full *Mafa-A* or *Mafa-B* genes or exons 2-4 of several of the classical class II genes (*Mafa-DRB, Mafa-DQA, Mafa-DQB, Mafa-DPA*) of cy0333 and three other MCM samples containing the M3 haplotype (data not shown). These gDNA amplicons cover ∼58 kb of the cy0333 hybrid ONT and PacBio MHC M3 haplotype assembly and differed at only two nucleotides, a single AG dinucleotide repeat within an ∼40 bp AG microsatellite sequence located in intron 2 of *Mafa-DRB*W049:01:01:01*. This equates to an estimated accuracy of 99.997% for the cy0333 hybrid assembly across the ∼58 kb examined by amplicon sequencing, but as observed directly in our assessment most of the nucleotide ambiguity occurs in challenging areas like microsatellite repeats and long homopolymers like poly-A or poly-T tracks across the full genomic MHC region.

### A potential recent expansion of MHC class I *AG*/*G*

The cy0333 MHC M3 haplotype of MCM shows evidence of a relatively recent segmental duplication event within the class I *Mafa-AG*/*Mafa-G* region (**Fig. 4**). There are two distinct instances of a virtually nucleotide-identical ∼47 kb block of sequence (differing at only 11 positions across 46,752 nucleotides) containing remarkably similar *Mafa-V*01:02* and *Mafa-G_pseudo1* genes plus three additional pseudogenes. Those blocks are followed by two minimally differentiated ∼48.5 kb blocks of sequence located just centromeric to each of the virtually nucleotide-identical *Mafa-G*/*Mafa-V* blocks. These neighboring blocks contain closely-related *Mafa-W* genes (*Mafa-W*01:11* and *Mafa-W*01:12*) and *Mafa-AG6* genes (*Mafa-AG6*04:05:03:03* and *Mafa-AG6*04:08:01:01*), plus three additional closely-related duplicated pseudogenes; these two blocks are ∼98.4% identical to each other (in contrast, each of the blocks containing *Mafa-AG6*04* genes were only ∼96.0% and ∼90.3% identical to the equivalent flanking blocks containing *Mafa-AG1*05:09:02:01* and *Mafa-AG2*01:05:01:01*, respectively). These highly conserved sequence blocks are not just an artifact of incorrect assembly because three independent ONT reads span both of the virtually identical *Mafa-G*/*Mafa-V* genes (**Fig. 4**). In fact, one of these ONT reads spans ∼546 kb and includes five of the six *Mafa-AG* genes of the M3 haplotype. A comparable recent duplication of the *Mamu-AG* region was previously observed in a well-characterized rhesus macaque MHC BAC library, with only eight nucleotide substitutions across an ∼80 kb sequence block (Daza-Vamenta et al. 2004).

**Figure 4.**
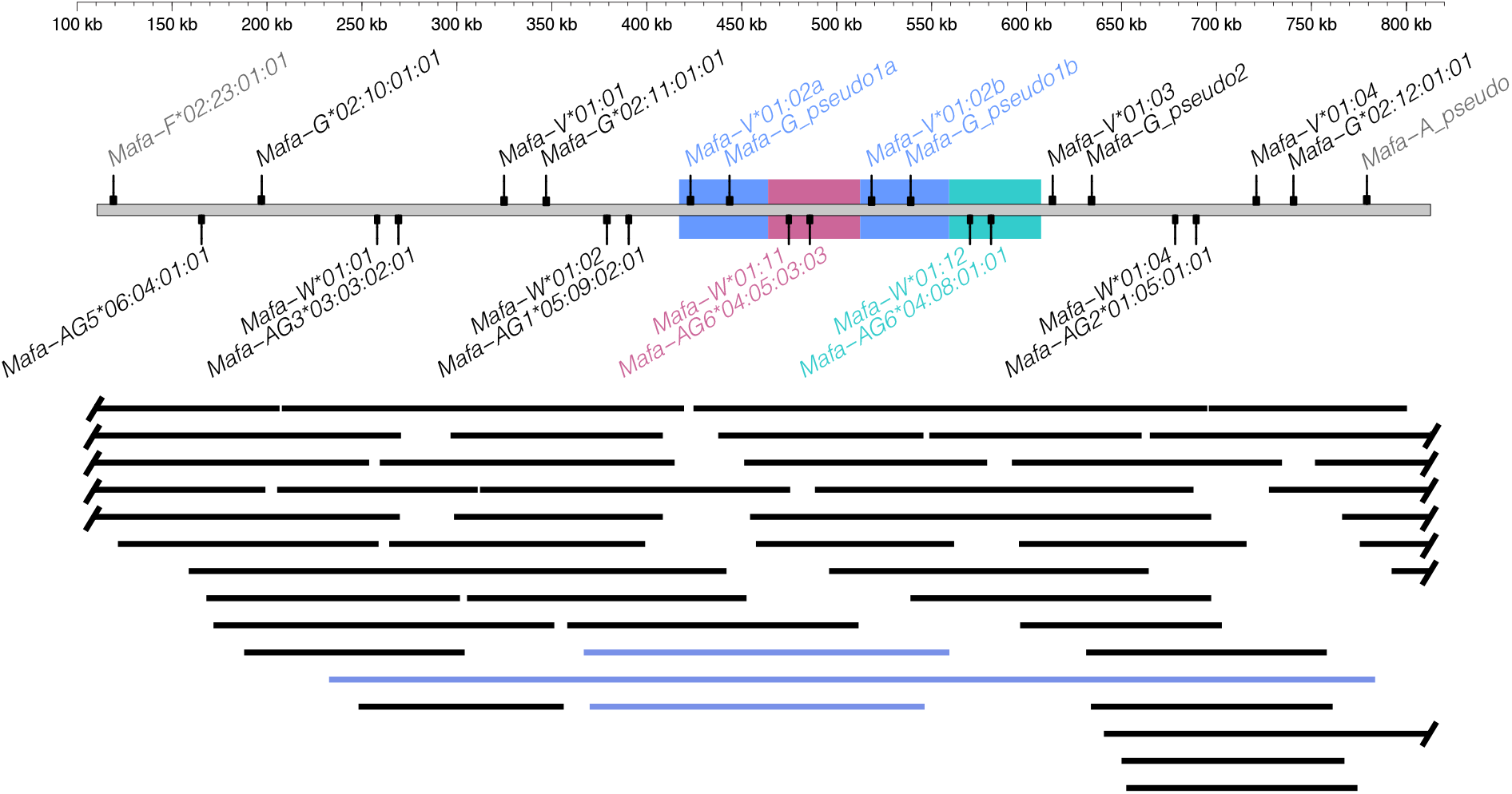
Gene content of the cy0333 MHC class I *AG/G* region. Representative genes and pseudogenes are shown on the gray bar. The blue boxes highlight a virtually nucleotide-identical ∼47 kb block of sequence located at two distinct positions within the class I *AG*/*G* region – this block contains remarkably similar *Mafa-V*01:02* and *Mafa-G_pseudo1* genes along with three additional pseudogenes not shown. The pink and green boxes highlight closely related (98.4%) ∼48.5 kb blocks of sequence located just centromeric to each of the virtually nucleotide-identical *Mafa-G*/*Mafa-V* blocks. The pink and green boxes contain closely-related *Mafa-W* genes (*Mafa-W*01:11* and *Mafa-W*01:12*, differing at 20 positions across 2,196 nucleotides) and closely-related *Mafa-AG6* genes (*Mafa-AG6*04:05:03:03* and *Mafa-AG6*04:08:01:01*, differing at 36 positions across 2,534 nucleotides), as well as three additional closely-related duplicated pseudogenes not shown. The position in kb relative to the start of the full genomic MHC region is shown across the top. The bottom portion shows the spans of individual ONT reads located within the class I *AG*/*G* region. The three ONT reads shown in blue span both copies of the essentially nucleotide-identical *Mafa-G*/*Mafa-V* genes.

### Specific duplicon structures associated with MHC class I B transcription levels in blood

The MHC class I B region contains 13 total *Mafa-B* genes associated with three different types of duplication blocks varying in the total pseudogene content of the area within and centromeric to each of those *Mafa-B* genes (**Fig. 5**). It is unknown how these variants in duplication blocks arose, if some of the duplication events were more localized and only included a portion of the surrounding pseudogenes. The first type of duplication block predates the divergence macaques and humans since it contains the full pseudogene content consistent with the *HLA-B* gene and its surrounding pseudogenes (*MIR6891, DHFRP2, RNU6-283P, FGFR3P1, ZDHHC20P2, HLA-S*). The second type of duplication block contains reduced pseudogene content, retaining only two or three of the surrounding pseudogenes, while the third type of duplication block only has a variant of the *MIR6891* microRNA located within intron 4 of each *Mafa-B* gene. The two major *Mafa-B* genes detected at high levels in steady-state whole blood cDNA (*Mafa-B*075:01:01:01* and *Mafa-B*011:01:01:01*) are both associated with the second type of duplication block with reduced surrounding pseudogenes (Budde et al. 2010; Wiseman et al. 2013). All five *Mafa-B* genes associated with full content blocks encode minor transcripts detected at low levels in steady-state whole blood cDNA. The sixth minor *Mafa-B* gene (*Mafa-B*079:01:01:01:01*) is the third type of duplication block with just the internal microRNA. The five *Mafa-B* pseudogenes containing atypical stop codons are all associated with reduced content duplication blocks; two are associated with the second type of duplication block (with just *MIR6891* and *DHFRP2*), and three are associated with the third type of duplication block (with just the internal microRNA *MIR6891*). Reduced surrounding pseudogene content roughly correlates to higher *Mafa-B* gene expression in steady-state whole blood unless the *Mafa-B* gene contains a premature or late stop codon rendering it unlikely to be translated into a traditionally functioning MHC class I protein. In comparison, full surrounding pseudogene content (consistent with *HLA-B*) roughly correlates to lower *Mafa-B* gene expression in steady-state whole blood. Notably, those surrounding pseudogenes are located centromeric to the anti-sense strand *Mafa-B* genes, positioning them upstream of the *Mafa-B* start codon.

**Figure 5.**
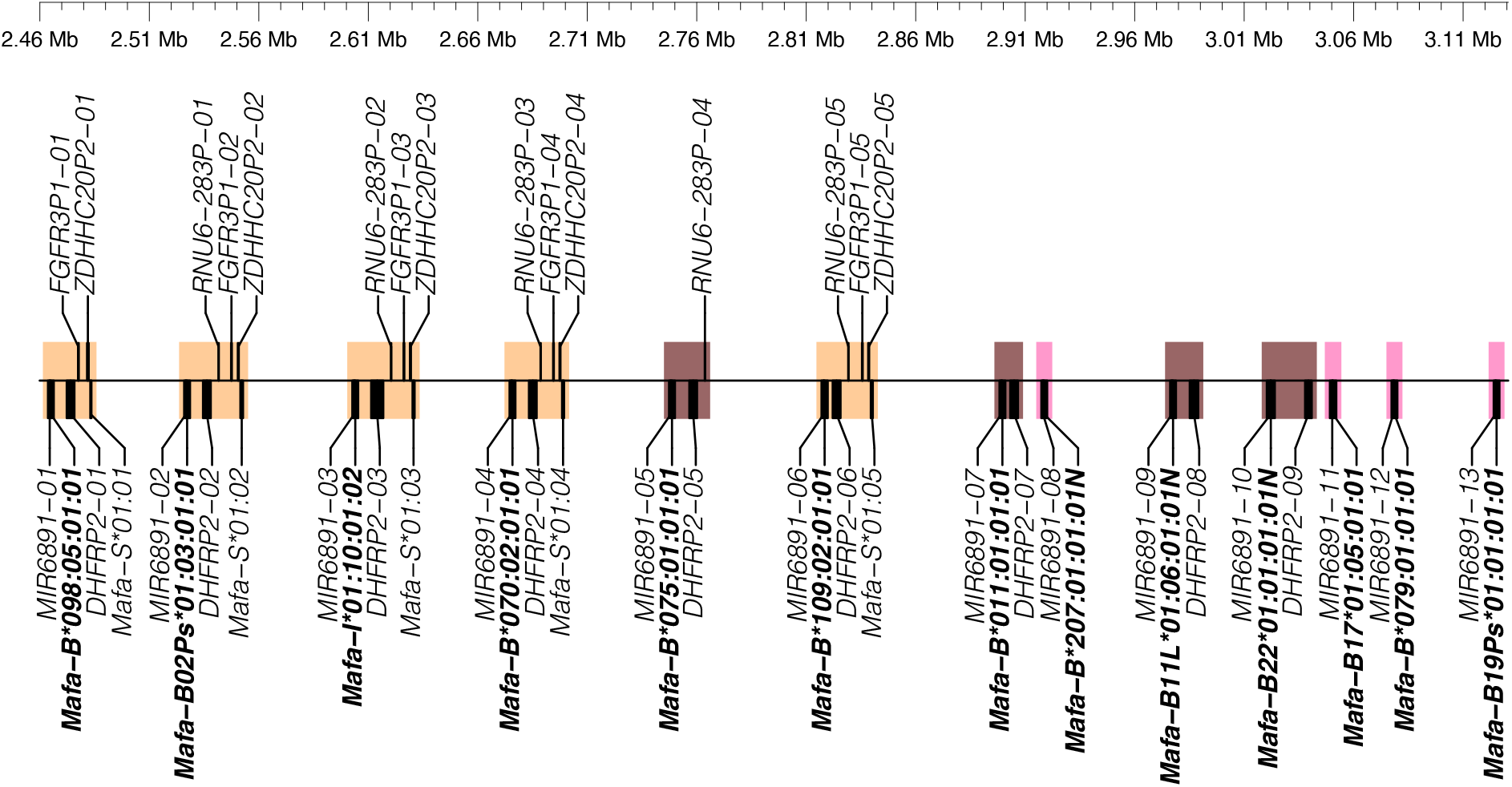
Gene content of the cy0333 MHC class I B region. Blocks of duplicated genes and pseudogenes are shaded. Orange shading indicates blocks containing essentially all six pseudogenes surrounding the classical class I *Mafa-B* gene, consistent with the pseudogenes surrounding the human *HLA-B* gene. Brown shading indicates blocks containing only two or three of the surrounding pseudogenes plus the *Mafa-B* gene. Pink shading indicates blocks containing only the *Mafa-B* gene and a microRNA located within intron 4 of the *Mafa-B* gene. *Mafa-B* allele names are displayed in bold. Position in Mb relative to the start of the full genomic MHC region is shown across the top.

None of the 13 *Mafa-B* gene duplication blocks contain pseudogene content consistent with the *HLA-C* gene and its surrounding pseudogenes (*USP8P1, RPL3P2, WASF5P, LINC02571*, and *LOC112267902*) (**Supplemental Table S1**). This is not unexpected, since *HLA-C* homologs have only been identified in great apes. It is thought that *HLA-C* and *HLA-B* developed from a common ancestral gene after apes and Old World monkeys like macaques diverged (Adams and Parham 2001; Fukami-Kobayashi et al. 2005).

## Discussion

### Utility of a highly accurate macaque MHC full genomic region assembly

The unambiguously assembled, contiguous, and highly accurate cy0333 MHC M3 haplotype of MCM presented here provides a valuable genomic region reference sequence. While the human HLA full genomic region serves reasonably well much of the time as a reference for extracting sequencing reads from a macaque data set for downstream *de novo* assembly, macaques may contain blocks of sequence within their MHC region that differ substantially enough from the human HLA region where no reads spanning that region will be extracted. We observed this directly when extracting the reads for error correction of our cy0333 MHC M3 region haplotype; a second round of mapping PacBio reads to our cy0333 assembly was required to extract additional reads to fill in a few areas left uncovered with the initial HLA mapping. Using the cy0333 MHC haplotype as the reference for future extractions of macaque sequence reads should substantially reduce this issue. The cy0333 MHC full genomic region haplotype is also suitable as a reference for mapping whole exome short reads from other macaque samples and is a helpful resource for designing amplicon primers, probes for target capture or CRISPR-Cas9 targeted enrichment as has recently been reported for macaque KIR gene clusters (Bruijnesteijn et al. 2021). The assembly of the full MHC genomic region from a homozygous macaque provides unequivocal confirmation that this cy0333 haplotype reference is an accurate ∼5.2 Mb end-to-end layout of the MHC M3 haplotype of MCM with no genes from other haplotypes mixed in, no genes expected on the M3 haplotype off on any secondary scaffolds, and no question of the validity of the assembly even with identical or highly similar sequence present more than once across the haplotype. This is especially valuable for assessing the gene content of the heavily duplicated MHC class I A and class I B regions. Additionally, the high accuracy of the PacBio HiFi reads generated for this assembly is valuable for distinguishing functional from non-functional genes, where single-nucleotide accuracy is vital for confirming true polymorphisms that introduce early stop codons into otherwise presumably functional genes.

### Classical MHC class I gene insights

When considering the wealth of class I genes carried on each MHC haplotype in macaques, we find it helpful to divide them into major versus minor allele lineages based on their steady-state RNA levels in whole blood. For the MHC M3 haplotype of MCM, these major alleles include *Mafa-A1*063:02:01:01, Mafa-B*075:01:01:01*, and *Mafa-B*011:01:01:01*. They appear to function like the *HLA-A* and *HLA-B* genes in humans and are responsible for the bulk of antigen presentation to CD8^+^ T cells (Loffredo et al. 2009; Budde et al. 2011). The number of major alleles varies between macaque MHC class I haplotypes; a single major A1 allele and two to four major B alleles per haplotype are commonly observed (Shortreed et al. 2020). The minor alleles of the M3 haplotype include *Mafa-A2*05:11:01:01, Mafa-A4*01:01:01:01, Mafa-B*070:02:01:01, Mafa-B*079:01:01:01, Mafa-B*098:05:01:01, Mafa-B*109:02:01:01*, and *Mafa-I*01:10:01:02*. Steady-state RNA levels for these minor alleles are reduced from 5-fold to more than 100-fold relative to the major alleles (Budde et al. 2011). Although they are present in variable numbers and combinations on every macaque MHC class I haplotype, the roles of these minor alleles in cellular immune responses remain elusive. In contrast to major allele lineages, the minor allele lineages tend to be distinguished by numerous allelic variants that are distributed across a wide variety of haplotypes (**Supplemental Table S2**). For example, the July 2022 IPD-MHC NHP database (Release 3.9.0.1 (2022-07) build 209) includes 95 and 85 distinct allelic variants of the *Mafa-A2*05* and *Mamu-A2*05* lineages, respectively. Likewise, characterization of a large breeding colony of Indonesian cynomolgus macaques revealed that nearly two-thirds (45/70) of the high-resolution *Mafa-A* haplotypes segregating in this population included one of at least 25 distinct *Mafa-A2*05* allelic variants (Shortreed et al. 2020). At least one member of the *A2*05* lineage (*Mamu-A2*05:01*) has been demonstrated to have the capacity to present simian immunodeficiency virus (SIV) epitopes (de Groot et al. 2017). Further studies will be required to untangle the full functions of these prevalent minor alleles such as *A2*05* in macaques.

Another long-standing puzzle that remains regarding the dramatic expansion of the MHC class I region described here is how macaques balance the maintenance of a sufficient repertoire of T-cell receptors to respond to a diversity of pathogens with negative selection to remove potentially autoreactive T cells (Daza-Vamenta et al. 2004). The collection of *Mafa-B* pseudogenes on the M3 haplotype provides a glimpse into this evolutionary balancing act. The loss of protein-coding potential for these class I pseudogenes has occurred due to an array of mechanisms. The first of these is exemplified with the *Mafa-B*207:01:01:01N* and *Mafa-B19Ps*01:01:01:01* alleles that have lost the appropriate in-frame ATG start codon due to single nucleotide variants. This appears to be an ongoing process of inactivation for the *Mafa-B*207* lineage since the internal in-frame start codon has also been disrupted for a second member of this lineage (*Mafa-B*207:03:01:01)*, but it is still intact for the closely related *Mafa-B*207:02:01:01* allele that retains the ability to encode a canonical class I protein. Unfortunately, the IPD-MHC NHP database currently lacks other members of the *Mafa-B19Ps* lineage besides the *Mafa-B19Ps*01:01:01:01* pseudogene. However, there is a *Mamu* homolog (accession # AB128842) of this *Mafa-B19Ps* pseudogene that was characterized as part of the complete rhesus macaque haplotype from the CHORI-250 BAC library (Daza-Vamenta et al. 2004; Shiina et al. 2006). Since both of these *B19Ps* pseudogenes carry identical start codon disruptions, this variant is likely to predate the speciation of cynomolgus and rhesus macaques more than 2 Myr ago. A second common mechanism of pseudogene formation is the introduction of a premature stop codon due to a single nucleotide variant in the coding region. The *Mafa-B22*01:01:01:01N* and *Mafa-B11L*01:06:01:01N* alleles of the M3 haplotype exemplify this mechanism with premature stop codons in exons 2 and 4, respectively. Another deviation from the traditional *MHC-B* open reading frame is observed in *Mafa-B17*01:05:01:01*. There is a single nucleotide deletion in the first codon of exon 6 of this allele, causing a frameshift through the rest of the open reading frame and loss of the typical *MHC-B* stop codon within exon 7. Since splicing sites compatible with the typical *MHC-B* exon 8 are retained, *Mafa-B17*01:05:01:01* would reach a stop codon 129 bp into the normal 3’ untranslated region and have a much longer cytoplasmic tail that is completely distinct from classical class I B proteins.

Transcriptional silencing appears to reflect yet another mechanism that may have led to the loss of functional class I protein production for the *Mafa-B02Ps*01:03:01:01* gene on the M3 haplotype. The IPD-MHC NHP database currently includes 17 *Mafa* and 24 *Mamu* full-length genomic alleles for this *B02Ps* lineage whose first representative alleles were deposited in 2010. Although cDNA sequences for this putative pseudogene lineage have yet to be detected despite exhaustive efforts by our group and other investigators, the mechanism of this transcriptional silencing phenomenon remains to be defined.

### Nonclassical MHC class I genes insights

Nonclassical MHC class I genes are often studied in the context of NK cell engagement. We characterized six distinct *Mafa-AG* alleles at the telomeric end of the MHC class I A region of cy0333. This region was organized into ∼40-80 kb genomic blocks with close sequence identity suggesting recent segmental duplication of the *Mafa-AG* locus. The *Mafa-AG* gene is believed to have functionally replaced the role of *HLA-G* in humans. Both genes are primarily expressed on the surface of invasive extravillous trophoblasts of the placenta and are thought to regulate the maternal immune response to pregnancy through the engagement of NK receptors. Indeed, rhesus macaque *Mamu-AG2*01:01* has been shown to have broad recognition by KIR genes *KIR3DL05, KIR3DL07, KIR3DS01, KIR3DS02, KIR3DSw07*, and *KIR3DSw09* (Nicholas et al. 2022; Anderson et al. 2022). Given the dramatic expansion of lineage II KIRs in macaques, iit is not surprising that the *MHC-AG* gene has also undergone tandem duplication events relative to the single copy *HLA-G* gene (de Groot et al. 2015). Our findings further implicate the close evolutionary relationship between the NK cell receptor and nonclassical *MHC-AG* gene clusters.

We also characterized three nonclassical class I *Mafa-E* sequences associated with the MHC M3 haplotype of MCM, two with complete traditional open reading frames and one non-coding pseudogene. The ability of MHC-E proteins to present MHC class I signal-sequence and pathogen-derived peptides to NK and CD8+ T cells is conserved between primate species (Wu et al. 2018). Cell surface expression of *MHC-E* can be induced by cytomegalovirus (CMV), and *MHC-E* has become a crucial target for CMV-based vaccine development (Tomasec et al. 2000; Voogd et al. 2022). Studies have demonstrated that 55% of rhesus macaques were completely protected against SIV challenge after vaccination with CMV/SIV vectors and that protection was mediated by *MHC-E*-restricted antigen recognition by CD8+ T cells (Hansen et al. 2011, 2013; Malouli et al. 2021). Genetic variation within *Mamu-E* genes has been demonstrated to be significantly associated with a lack of CMV/SIV vaccine protection in this model (Brochu et al. 2022). Given this association, the simplified genetics of MCM may serve as a better model system to dissect the genetic basis of protection in CMV-based vaccination studies. Furthermore, examination of transposable elements, untranslated region sequences, and intronic polymorphisms from multiple MHC genomic haplotypes within the MCM population may provide a clearer image of the genetic determinants of vaccine-mediated protection.

### Caveats and future directions

Since the conception of this manuscript, a new ONT base caller, Remora, with marginally higher single nucleotide accuracy was commercially released (https://github.com/nanoporetech/remora). Though higher accuracy base calls will theoretically benefit the ONT assembly process, we are confident that our hybrid assembly does not require the marginal gains of Remora due to the end-to-end PacBio HiFi coverage used in error correction. We assessed the accuracy of MHC class I and class II genes in our hybrid cy0333 assembly versus gDNA amplicon sequences and found 99.997% concordance, with the only variance occurring within a short tandem repeat. While single nucleotide accuracy is improved with Remora, it remains to be seen if similar accuracy gains are achieved in difficult areas like repetitive motifs.

Alternate assembly algorithms such as FLYE and Wengan D can produce highly accurate assemblies using multiple sequencing technologies (Kolmogorov et al. 2019; Di Genova et al. 2021). We used both FLYE and Wengan D in our initial assembly attempts and found both incapable of assembling the highly-duplicated *MHC-AG* region. In contrast, Canu was capable of properly untangling the *MHC-AG* region, which we confirmed by manually inspecting ONT reads spanning the neighboring, near-identical *Mafa-AG* genomic blocks. Therefore, for our purposes, Canu seems to be the most capable of handling highly repetitive genomic regions.

The approach to *de novo* MHC full genomic region assembly through whole genome sequencing is admittedly cost-prohibitive for large-scale MHC haplotype characterization of large cohorts of animals. Recently, Cas9 enrichment was used to create haplotype-phased genomic assemblies of rhesus macaque KIR regions (Bruijnesteijn et al. 2021). A similar approach can most likely be applied towards MHC enrichment; however, we forwent region-specific enrichment in favor of unbiased, whole genome sequencing in order to create assemblies of multiple regions from the same animal. Sequencing all of the immune-important gene clusters (spread across different chromosomes) from the same animal might point to insights into specific protein interactions that would be lost when examining each genomic region independently from different samples.

The method presented here for assembling the cy0333 MHC M3 haplotype of MCM from ultra-long ONT and highly accurate PacBio HiFi sequencing reads can be directly applied to resolving the full MHC genomic regions of the remaining six haplotypes in the MCM population, provided samples are obtained from additional MHC-homozygous individuals. It can also be extended to the characterization of sire-dam-offspring trios from additional nonhuman primate species where sequence reads can be binned to assemble appropriately phased haplotypes per animal to avoid chimeric MHC assemblies. With additional unambiguous full macaque MHC regions solved, it may be possible to assess any potential genomic cues for major versus minor differential expression of the MHC class I genes in steady-state RNA from whole blood. Finally, this method can also potentially be extended to assembly and annotation of other immune-important genomic regions of macaques such as the leukocyte receptor complex and KIRs, natural killer complex, Fc gamma receptors, T cell receptors, and immunoglobulin gene clusters. Together with the restricted genetic diversity of the MCM population, the development of these new genomic resources will provide exceptional opportunities to improve the design and reproducibility of a wide variety of biomedical research.

## Methods

### Samples

Splenocytes and peripheral blood mononuclear cells (PBMC) were obtained from a Mauritian-origin cynomolgus macaque (cy0333) housed at the Wisconsin National Primate Research Center. This macaque was previously genotyped as M3 homozygous in the MHC region (Ericsen et al. 2014). Sampling was performed following protocols approved by the University of Wisconsin-Madison Institutional Animal Care and Use Committee and in accordance with the regulations and guidelines outlined in the Animal Welfare Act, the Guide for the Care and Use of Laboratory Animals, and the Weatherall report (Weatherall 2006).

### High molecular weight DNA extraction for Oxford Nanopore sequencing

DNA was extracted from frozen splenocytes or PBMC using slight modifications of a previously described ultra-long DNA extraction protocol (Jain et al. 2018). Per extraction, 2 × 10^7^ cells were thawed and spun at 300 x *g* for 10 min. Pelleted cells were resuspended in 200 μL PBS by pipette mixing. Resuspended cells plus 10 mL TLB (100 mM NaCl, 10 mM Tris-Cl pH 8.0, 25 mM EDTA pH 8.0, 0.5% (w/v) SDS, 20 μg/mL Qiagen RNase A, H_2_O) were combined in a 15 mL conical tube and vortexed at full speed for 5 sec. Samples were incubated at 37°C for 1 hr. Qiagen Proteinase K (200 μg/mL) was added and mixed by slow end-over-end inversion three times. Samples were incubated at 50°C for 2 hrs with 3x end-over-end inversion every 30 min. The lysate was slowly pipetted into two phase-lock 15 mL conical tubes (prepared by adding ∼2 mL autoclaved high-vacuum silicone grease into 15 mL conical tubes and spinning max speed for 1 min to move the grease to the bottom of the tube), adding 5 mL sample to each tube. To each phase-lock tube of lysate, 2.5 mL buffer-saturated phenol and 2.5 mL chloroform were added before rotational mixing at 20 rpm for 10 min. Tubes were then spun down at 4,000 rpm for 10 min in an Eppendorf 5810R centrifuge with a swing-bucket rotor. After centrifugation, the aqueous phase above the silicone grease from each conical tube was poured into a new phase-lock 15 mL conical tube. The addition of phenol and chloroform, rotational mixing, and centrifugation was repeated with the second set of phase-lock tubes. The aqueous phases from both phase-lock tubes were slowly poured into a single 50 mL conical tube before adding 4 mL of ammonium acetate (5 M) and 30 mL of ice-cold 100% ethanol. The mixture was incubated at room temperature while the DNA visibly precipitated. Once the DNA rose to the surface of the mixture, a glass capillary hook was used to retrieve the DNA. The hooked DNA was dipped into 70% ethanol to tighten the DNA into an opaque pellet on the hook, then carefully worked off of the hook into an Eppendorf DNA LoBind 1.5 mL tube. To wash the DNA pellet, 1 mL of 70% ethanol was added to the tube before spinning down at 10,000 x *g* for 1 min, then removing as much ethanol as possible. This process was repeated for a second 70% ethanol wash, and any remaining ethanol was evaporated off during a 15 min incubation at room temperature. Finally, 100 μL of elution buffer (10 mM Tris-Cl pH 8.0, H_2_O) was added before incubating the DNA at 4°C for at least two days to resuspend fully. A total of eleven ultra-long DNA isolations were performed to obtain enough product for ONT sequencing.

### High molecular weight DNA extraction for PacBio HiFi sequencing

DNA was extracted from PBMC using New England Biolab’s Monarch High Molecular Weight DNA Extraction Kit for Cells and Blood following the manufacturer’s protocol. Briefly, 5 × 10^6^ PBMC were pelleted by centrifugation for 3 min at 1,000 x *g*. 150 μL of Nuclei Prep Solution was added to the resuspended pellet and pipetted 10 times up and down to mix. 150 μL of Nuclei Lysis Solution was added and inverted 10 times to mix. Each sample was incubated at 56°C for 10 min in a thermal mixer at 2,000 rpm. Following incubation, 75 μL of precipitation buffer was added and the solution was inverted 10x to mix. Two DNA capture beads were added to each sample, followed by 275 μL of isopropanol. Samples were inverted on a vertical rotating mixer at 10 rpm for 8 min. The liquid was pipetted from the samples, avoiding gDNA wrapped around the capture beads. Two washing steps were performed by adding 500 μL of DNA wash buffer, inverting 3 times, then carefully removing the wash buffer by pipetting while avoiding the DNA capture beads. Following washes, the beads were poured into a supplied bead retainer and pulse spun for ∼1 sec. The separated glass beads, along with 100 μL of elution buffer, were added to a 2 mL microfuge tube and incubated for 5 min at 56°C in a thermal mixer at 300 rpm. After incubation, the eluate was separated from DNA capture beads using the supplied bead retainer and poured into an Eppendorf DNA LoBind 1.5 mL tube. The bead retainer and 1.5 mL tube were centrifuged for 1 min at 12,000 x *g* to separate eluate from DNA capture beads. The final eluate with DNA was stored at 4°C until library preparation.

### Library preparation for Oxford Nanopore sequencing

Ultra-long DNA was prepared for sequencing on ONT R9.4 MinION flow cells using the SQK-RAD004 Rapid Sequencing Kit with modifications of a previously described library preparation optimized for the RAD004 sequencing kit (Jain et al. 2018). A 16 μL aliquot of DNA was pipetted as slowly as possible, using a cut-off P20 pipette tip, into a 0.2 mL PCR tube. 1 μL of that DNA aliquot was used for quantification using the ThermoFisher Scientific Qubit dsDNA BR Assay Kit. Due to the heterogeneity of ultra-long DNA, concentrations for the same DNA isolation were highly variable. Qubit concentrations ranged from 43.6 ng/μL to 450 ng/μL per quantified aliquot, with an average concentration of 231.1 ng/μL across all aliquots. To the remaining 15 μL of DNA in the PCR tube, 1.5 μL fragmentation mix (FRA) and 3.5 μL elution buffer were added, then pipetted up and down as slowly as possible 8x to mix using the cut-off P20 pipette tip. The sample mixture was thermal cycled to 30°C for 1 min, 80°C for 1 min, then a 4°C hold to perform the fragmentation. 1 μL rapid adapter (RAP) was added, then pipetted up and down as slowly as possible 8x using the cut-off P20 pipette tip to mix. The DNA mixture was incubated at room temperature while the flow cell was primed.

Flow cell priming was performed by adding 30 μL of flush tether (FLT) to a tube of flush buffer (FB), then vortexing the FB tube and briefly spinning it down. A flow cell check was performed, then a tiny volume of storage buffer was removed from the flow cell and 800 μL of flush mix was added through the priming port following the manufacturer’s instructions. After 5 min, the SpotON port was opened and another 200 μL of flush mix was added through the priming port, watching for a small amount of liquid to bead up through the SpotON port. Returning to the DNA mixture, 34 μL sequencing buffer (SQB) and 20 μL H_2_O were added and slowly pipetted up and down 5x with a wide-bore P200 pipette tip to mix. The final DNA mixture was then loaded dropwise into the SpotON port for diffusion loading of the MinION flow cell.

### MinION sequencing and base calling

Sixty R9.4 (FLO-MIN106) MinION flow cells of ultra-long DNA were sequenced according to ONT guidelines using their MinKNOW software to control each sequencing run. Different versions of MinKNOW were used throughout the course of the study as updates were released; specific versions are recorded in the metadata in the fast5 files for each run. Base calling was performed using Bonito version 0.3.8 on A100 GPU hardware running CUDA 11.2.

### PacBio Sequel II sequencing

High molecular weight DNA was provided to the University of Wisconsin-Madison Biotechnology Center DNA Sequencing Facility. The quality of the extracted DNA was measured on a ThermoFisher Scientific NanoDrop One instrument. Concentrations, 260/230 ratios, and 260/280 ratios were logged. Quantification of the extracted DNA was measured using the ThermoFisher Scientific Qubit dsDNA High Sensitivity kit. Samples were diluted before running on the Agilent FemtoPulse System to assess DNA sizing and quality. A PacBio HiFi library was prepared according to PN 101-853-100 Version 03 (Pacific Biosciences). Modifications include shearing with Covaris gTUBEs and size selection with Sage Sciences BluePippin. Library quality was assessed using the Agilent FemtoPulse System. The library was quantified using the Qubit dsDNA High Sensitivity kit. The library was sequenced on a PacBio Sequel II instrument using Sequel Polymerase Binding Kit 2.2 at the University of Wisconsin-Madison Biotechnology Center DNA Sequencing Facility. Raw sequencing data was converted to circular consensus sequencing (CCS) fastq using SMRTLink version 8.0.

### Hybrid *de novo* assembly of the MHC region

ONT and PacBio HiFi fastq reads were mapped against the human reference genome GRCh38 (GCA_000001405) using minimap2 version 2.17 (Li 2018) using the ‘-ax map-ont’ and ‘-ax map-hifi’ flags, respectively. Reads mapping to coordinates chr6:29,400,000-33,525,000 (the coordinates for the genes bounding the full genomic HLA region, *GABBR1* through *KIFC1* (Shiina et al. 2017), plus ∼200 kb of sequence on both ends) were extracted using samtools version 1.11 and converted back to fastq using bbtool’s reformat.sh (https://sourceforge.net/projects/bbmap/). Extracted ONT and PacBio reads were assembled separately using FLYE version 2.7 using the ‘--genome-size 5m’ and ‘--nano-corr’ or ‘--pacbio-hifi’ flags. After assembling, the ONT-only and PacBio-only contigs were merged using quickmerge version 0.3 (https://github.com/mahulchak/quickmerge) to produce a hybrid assembly. This assembly produced seven total contigs, the longest being ∼5.4 Mb. Upon closer investigation of the longest hybrid contig, we noticed a break within the *Mafa-AG* region. Based on our previous experience with the MHC regions of macaques, we were aware that the *Mafa-AG* duplicons within the MHC class I A region are challenging to resolve due to the potential for large blocks of highly similar sequence (Boyson et al. 1999; Daza-Vamenta et al. 2004; Shiina et al. 2006). We therefore decided that FLYE by itself may not be capable of properly assembling this highly repetitive genomic region and that Canu may serve as a better alternative for this region. To attempt to better focus a second *de novo* assembly on the region where the previous assembly broke, reads mapping to coordinates chr6:29,400,000-30,033,000 (the coordinates for the genes bounding the HLA class I A region, *GABBR1* through *POLR1H* (Shiina et al. 2017), plus flanking sequence) were extracted and assembled separately using Canu version 2.2 with the ‘-genomeSize=1m’ and ‘-nanopore’ or ‘-pacbio-hifi’ flags, respectively. ONT and PacBio contigs were then merged using quickmerge version 0.3. The second assembly produced a single ∼1.5 Mb contig. A comparison between the initial full genomic MHC region assembly and the secondary class I A-focused assembly showed excellent concordance in the areas of overlap; the two assemblies were then manually combined at the start of the predicted *Mafa-J* gene to create one scaffold 5,227,476 bp in total length, spanning from *GABBR1* through *KIFC1*. Metrics for numbers of ONT and PacBio reads at or above various lengths for both the MHC region and the whole genome are shown in **Supplemental Table S3** and read coverage for ONT and PacBio across the full genomic MHC region are shown in **Supplemental Fig. S1**.

### PacBio HiFi polishing

First-round error correction was performed using the extracted PacBio HiFi reads that aligned to human coordinates chr6:29,400,000-33,525,000 with Geneious Prime v 2022.1.1 (https://www.geneious.com/). The Geneious mapper was used with custom sensitivity and iteration up to 10 times for fine-tuning. Several advanced custom settings were enabled on Geneious: map multiple best matches to none, trim paired read overhangs, only map paired reads which both map, allow a maximum of 10% gaps per read with maximum gap size of 15, word length of 18 and index word length of 13, ignore words repeated more than 12 times, maximum of 5% mismatches per read, maximum ambiguity of 4, and accurately map reads with errors to repeat regions. A consensus sequence for that mapping was generated in Geneious using a 0% (majority) threshold, assigning total quality and calling the reference (the hybrid scaffold) if no coverage, and calling the reference if coverage was less than 5 reads. The resulting consensus sequence was thus corrected to correspond to higher-accuracy PacBio HiFi reads at any positions where the PacBio data differed from the hybrid assembly provided the PacBio coverage was greater than five reads. A few areas of the hybrid assembly lacked any PacBio coverage and thus remained uncorrected. These regions presumably corresponded to areas of the macaque MHC that differed substantially from the human HLA region. To address these areas, we performed a second round of PacBio mapping using minimap2 and extracted reads that aligned to the first-round error corrected hybrid assembly. This new set of PacBio reads was then used to error correct a second time and fill in gaps left by the HLA-extracted PacBio reads. The final mapping resulted in end-to-end, >5x PacBio coverage across the entire ∼5.2 Mb hybrid assembly (**Supplemental Fig. S1**).

### Gene annotation

We used the sequence comparison tool Exonerate version 2.4.0 using the ‘est2genome’ mapping model. Exonerate was run recursively using three different sequence lists: human gene and CDS annotations from GRCh38.p12, rhesus macaque MHC class I region CDS annotations from CHORI-250 BAC library from GenBank sequence AB128049 (Shiina et al. 2006), and cynomolgus macaque CDS sequences from the IPD-MHC NHP release 3.2.0.0 (Maccari et al. 2017). Exonerate results for each sequence list were filtered to matches of >95%, and those annotation tracks were loaded onto the polished cy0333 sequence scaffold in Geneious. Annotations were manually examined and curated to retain only a single gene annotation per locus, plus CDS, ncRNA, and exon annotations as appropriate. All annotated genes were individually compared against all available human and rhesus macaque orthologs to confirm a proper annotation. In the absence of any transcription data, gene and CDS annotations spanned from start codon to stop codon for presumed transcribed genes and corresponded to human and rhesus macaque gene annotations for pseudogenes and noncoding RNAs. Gene names were assigned based on human and rhesus macaque orthologs.

### Figure generation

Gene, pseudogene, and ncRNA annotations were created from gff3 files exported from Geneious. Genomic annotation maps were generated from exported gff3 files using the KaryoploteR package (Gel and Serra 2017) and RStudio (https://www.rstudio.com). Additional formatting and editing to improve readability of gene symbols were performed using Adobe Illustrator.

### Oxford Nanopore-based whole genome reference

While this report focuses on the full genomic MHC region, there was no feasible and economical way to sequence ultra-long reads corresponding to just the MHC selectively. Therefore, we sequenced the full cy0333 MCM genome with ultra-long ONT reads. Macaca_fascicularis_6.0 (GCA_011100615) was generated from ∼53.6x genome coverage ONT reads with corresponding BioNano optical mapping for assembly guidance. A well-characterized full genomic MHC region, along with a similarly detailed full genomic KIR gene region, are included within that Macaca_fascicularis_6.0 assembly. That assembly was error corrected with Illumina short-read data, so accuracy is not as high as the full genomic region MHC hybrid ONT and PacBio assembly presented here (OP204634).

## Data Access

Raw whole genome Oxford Nanopore reads were submitted to the National Center for Biotechnology Information (NCBI) Sequence Read Archive (SRA) under accession number SRX15982296. Raw whole genome PacBio HiFi reads were submitted to NCBI SRA under accession number SRX15982297. The final full ∼5.2 Mb annotated MHC genomic region sequence was submitted to NCBI GenBank under accession number OP204634. Classical and nonclassical MHC class I and class II genes were submitted to NCBI GenBank and the IPD-MHC NHP for official nomenclature under GenBank accession numbers MW679584-MW679656 and ON736434-ON736437 (Maccari et al. 2017; de Groot et al. 2020). All custom scripts are also available from https://go.wisc.edu/svvlra.

## Acknowledgments

The authors thank Miranda Stauss for assistance with PacBio MHC amplicon analysis and Nicholas Minor for assistance with KaryoploteR and RStudio. We are also grateful for the expert assistance of Natasja de Groot in providing the official IPD-MHC NHP allele nomenclature for the MHC class I and class II alleles reported here. We also thank R. Alan Harris, Muthuswamy Raveendran, and Jeffrey Rogers for their whole genome assembly efforts to generate Macaca_fascicularis_6.0. This work was supported through contracts HHSN272201600007C and 75N93021C00006 from the National Institute of Allergy and Infectious Diseases of the NIH. The authors utilized the University of Wisconsin-Madison Biotechnology Center’s DNA Sequencing Facility (Research Resource Identifier – RRID:SCR_017759) to generate and sequence PacBio Sequel II HiFi libraries. This work was also partly supported by the Office of Research Infrastructure Programs P51OD011106 awarded to the Wisconsin National Primate Research Center at the University of Wisconsin-Madison, and conducted in part at a facility constructed with support from Research Facilities Improvement Program grants RR15459-01 and RR020141-01.

