## Supplemental Fig S1 for "Complete sequencing of a cynomolgus macaque major histocompatibility complex haplotype"

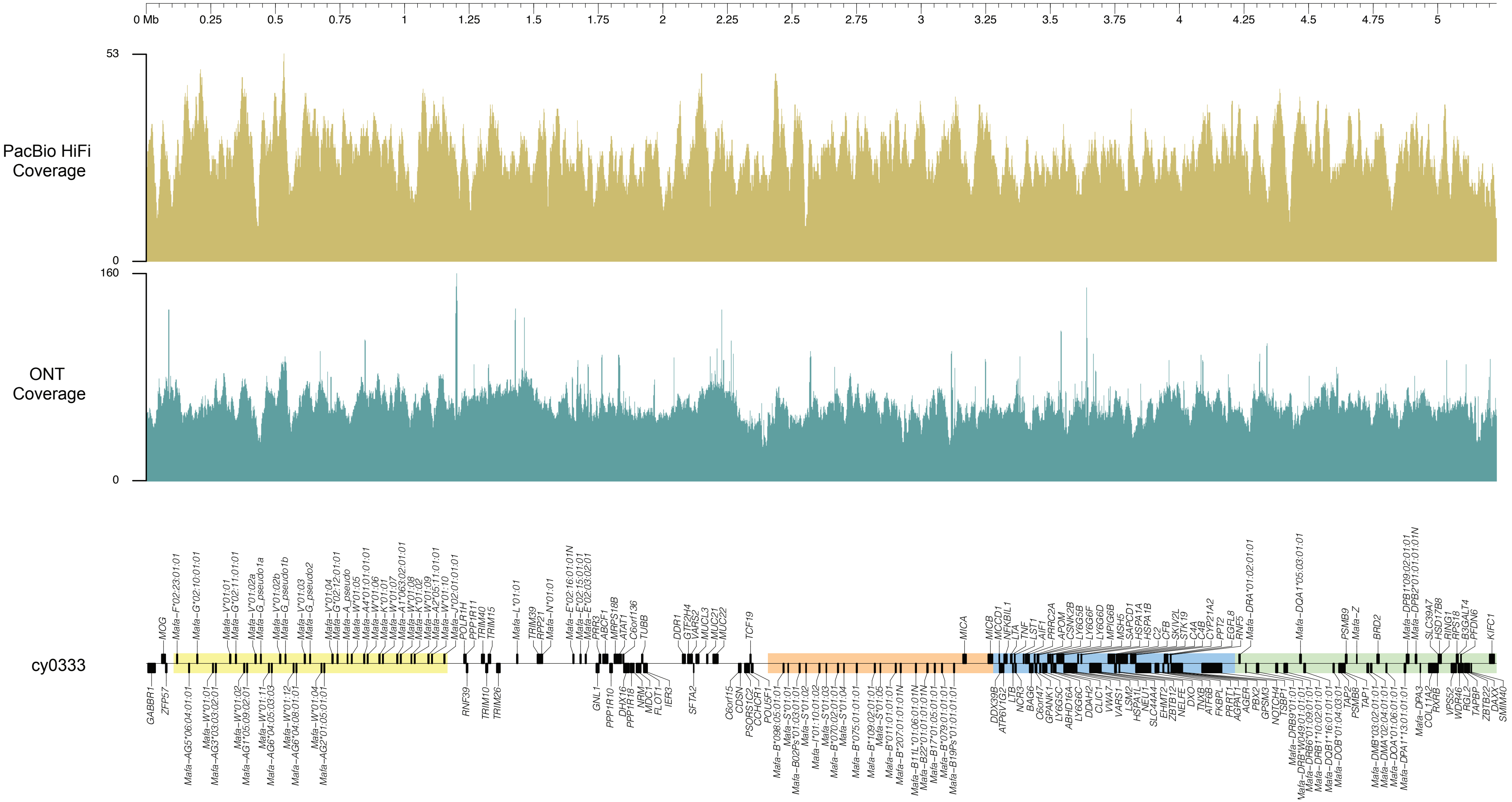


**Supplemental Figure S1.** Oxford Nanopore and PacBio HiFi read coverage for cy0333 MHC assembly. Coverage was determined by mapping extracted fastq reads against the error-corrected, merged assembly using minimap2 and plots were generated using R. Length is depicted in Mb. Representative assembly annotations are depicted on the bottom track. Coding regions are depicted by black boxes on top of or below the assembly. The class I A, class I B, class III, and class II regions are depicted by yellow, orange, blue, and green highlighting, respectively.
